# OpTiles: An R Package for Adaptive Tiling and Methylation Variability Profiling

**DOI:** 10.1101/2025.09.04.674166

**Authors:** Giorgia Migliaccio, Lena Möbus, Giusy del Giudice, Jack Morikka, Antonio Federico, Angela Serra, Dario Greco

## Abstract

**Summary:** OpTiles is an R package that dynamically defines tiling windows based on the distribution of sequenced CpGs, addressing the limitations of traditional fixed-tiling approaches in targeted methylation datasets. By integrating CpG density with intra-region methylation variability, it provides a reliability metric and extended functionality for annotating, prioritizing, and interpreting complex methylation data.

**Availability:** OpTiles is implemented in R and source code is freely available at https://github.com/fhaive/OpTiles. Data are available on Zenodo at https://doi.org/10.5281/zenodo.16961293

## Introduction

The advent of next-generation sequencing technologies has significantly advanced epigenomic research, enabling genome-wide profiling of DNA methylation at single-CpG resolution (Lister et al. 2009, Plongthongkum, Diep and Zhang 2014). However, despite this high-resolution capability, it remains standard practice to aggregate neighbouring CpG sites into larger regions to capture broader methylation patterns and extract biologically meaningful signals(Akalin et al. 2012, Hansen, Langmead and Irizarry 2012, Li et al. 2013, Gaspar and Hart 2017, Piao et al. 2021). Region-based strategies have been developed to define regions from CpG-level patterns. Several methods, such as DSS (Feng, Conneely and Wu 2014) and bsseq (Hansen, Langmead and Irizarry 2012) model methylation as a continuous signal across the genome, using smoothing or hierarchical frameworks to capture spatially coherent methylation patterns across neighbouring CpGs. Others, like metilene (Jühling et al. 2016) and DMRcate (Peters et al. 2015), identify and cluster differentially methylated CpGs based on proximity and methylation status. Other approaches as DMAP (Stockwell et al. 2014) define regions based on the actual DNA fragments obtained during library preparation, effectively anchoring methylation summaries to experimental fragment boundaries. The approaches described here represent only a subset of the many strategies reviewed in the literature (Piao et al. 2021, Chen, Lin and Fann 2016, Shafi et al. 2018), where a variety of methodologies have been proposed to aggregate CpG sites into regions. However, no consensus has been reached on what constitutes the optimal strategy.

As noted by Shafi and colleagues (Shafi et al. 2018), one of the most widely cited approaches is implemented in methylKit (Akalin et al., 2012), which uses a tiling-window strategy to divide the genome into fixed-size segments and summarize the methylation signals of CpGs within each of them. While this approach offers systematic coverage it assumes a uniform distribution of CpGs and consistent sequencing coverage which limits its suitability for targeted approaches like Reduced Representation Bisulfite Sequencing (RRBS), where the CpG distribution is biased towards CpG-rich regions within the genome. As a result, fixed tiles may span poorly covered sites, potentially diluting signals and introducing noise. Filtering tiles by CpG density is therefore common practice, but when region boundaries are not aligned with the distribution of sequenced data, this filtering can inadvertently discard regions that still hold valuable information. Furthermore, the summarization approach used in methylKit (Akalin et al., 2012) reduces each region to a single average methylation value, without accounting for intra-region variability. Some methods treat CpG methylation consistency as a requirement for defining regions (Balaramane et al. 2024, Shi et al. 2021, Hansen, Langmead and Irizarry 2012), instead in the most straightforward genome-tiling approach methylation consistency within regions is overall ignored. Although colocalised CpGs are generally expected to exhibit coordinated methylation patterns (Eckhardt et al. 2006, Affinito et al. 2020), heterogeneity in methylation states is frequently observed (Elliott et al. 2015). Such variability can be influenced by technical factors, such as PCR bias, sequencing coverage, different read lengths or stochastic erosion (Scherer et al. 2020). PCR amplification may preferentially enrich molecules with particular base compositions or methylation states, while incomplete bisulfite conversion, uneven sequencing coverage, or readmapping errors can introduce spurious heterogeneity across CpGs (Scherer et al. 2020). However, methylation heterogeneity may also reflect variation in cell-type composition, allele or stand specific methylation, imprinted regions, or even transcriptional heterogeneity (Olova et al. 2018, Shi et al. 2021). By incorporating variability metrics into fixed- tiles, OpTiles adds this missing layer of context by enabling the identification of regions with high intra-region variability and guiding decisions on whether to summarize a genomic locus as a region or retain the finer resolution of individual analysis.

In this study, we introduce OpTiles, an R package built as a modular extension to the methylKit workflow (Akalin et al. 2012) which leverages its well-established tiling-window framework to enhance methylation region definition. While designed to integrate seamlessly with methylKit, OpTiles’ core functions, such as optimization of tiles definition, intra-region variability assessment and annotations functions, require only genomic coordinates and CpG-level beta values as input, enabling the use outside the methylKit environment. Its core functions are modular and can operate independently, while auxiliary functions are also provided to convert data into methylKit format for running parts of its standard workflow (e.g. loading data or differential methylation analysis), facilitating direct integration into existing pipelines. The manual (on https://github.com/fhaive/OpTiles) explains in detail how the function works and what inputs are needed. Rather than applying uniform, fixed-size tiling across the genome, OpTiles refines the tiles *post hoc* by repositioning them based on the distribution of sequenced CpGs, ensuring tiles to be anchored in sequenced CpGs. Additionally, OpTiles provides functions to assess intra-region methylation variability, an often overlooked dimension, that can help users identify high variable regions, either technically unstable or biologically heterogeneous, supporting more informed filtering and prioritization decisions. Beyond tile optimization and variability assessment, OpTiles maps user-defined regions to genomic annotations and provides overlap metrics that consider both region size and CpG content. Together, these functions support a more informed and flexible interpretation of DNA methylation landscapes, complementing existing pipelines without altering their core analytical logic.

## Optimization of tiles definition

OpTiles refines the definition of tiling window with a data-driven adaptive binning: starting from the tiling window genomic locations, OpTiles computes the number of CpGs within each tile (with *map_cpg_to_regions* function) and identifies pairs of consecutive tiles defined as directly adjacent, non-overlapping pairs. For each pair of candidate tiles, OpTiles maps all CpG positions and calculates two distances (Figure 1A): (1) AD is the distance between the first CpG of the first tile and the last CpG of the second; (2) BC is the distance between the last CpG of the first tile and the first CpG of the adjacent one. If CpGs are tightly clustered (AD < tile size), the tiles are merged into a single region spanning all CpGs from both tiles. If AD > tile size but BC < tile size, indicating a narrow gap between CpG clusters, OpTiles scans the span to generate fixed size windows and selects the one with the highest CpG count, retaining the most downstream in case of a tie. If neither AD nor BC meet these criteria, the tiles remain unchanged. This optimization is performed using the *merging_consecutive_regions* function, which takes as input the original CpGs genomic location matrix, the location of the tiles, and the regions length.

**Figure 1:**
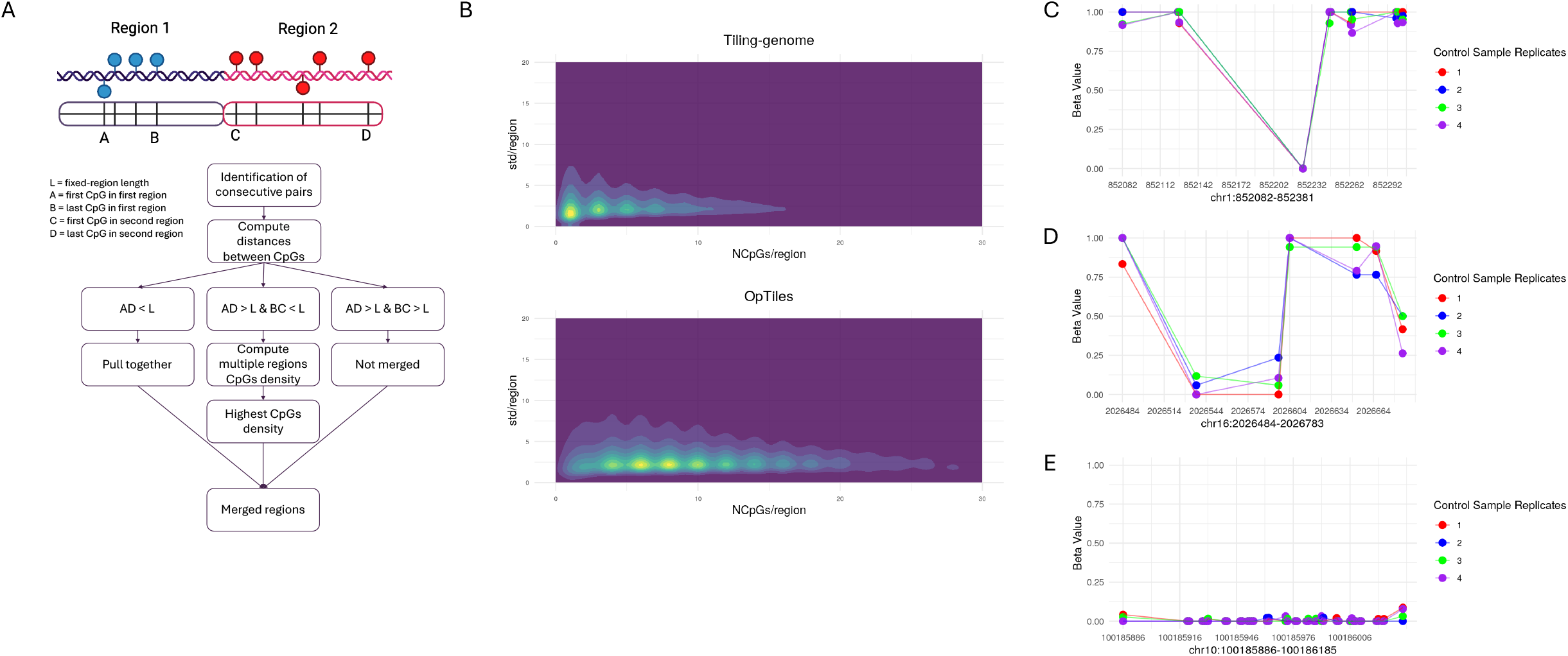
(A) Schematic overview of the workflow for optimizing tile definitions. (B) Comparison of average CpG density per region before and after applying the optimization strategy. (C), (D), (E) Intra-region methylation variability of three selected genomic region.

To illustrate the impact of our merging strategy, we analysed an RRBS dataset from THP-1–derived macrophages (data available on Zenodo https://doi.org/10.5281/zenodo.16961293). THP-1 cells (ATCC TIB-202, USA) were cultured in RPMI-1640 (Gibco, USA) supplemented with 10 % FBS (Gibco, USA) (culture media). Cells were cultured in 75 cm^2^ flasks at a density < 1 × 10^6^ cells/ml. Cells were differentiated in 12 well plates, with 127,000 cells/cm^2^, in the culture media supplemented with 30.9 ng/ml of phorbol 12-myristate 13-acetate (PMA) (Sigma-Aldrich, USA) for 48 hours, followed by a 72-hour recovery period in standard culture medium. The preprocessing of the RRBS data has been done using Bismark suite (Krueger and Andrews 2011) and methylKit pipeline (Akalin et al. 2012) (script available on git), and regions produced by the standard methylKit tiling were compared with those from our optimized approach. As shown in Figure 1B, CpG density is higher in the optimized tiles compared with the standard tiles. This improvement is particularly useful given that many pipelines apply minimum-CpG filters to remove low-information regions (Akalin et al. 2012, Li et al. 2013, Dolzhenko and Smith 2014, Piao et al. 2021). For example, applying a >5 CpG filter to our dataset removed 45.6% of standard tiles but only 22.3% of optimized tiles (Supplementary Table 1). By increasing CpG density per region, OpTiles preserves potentially informative regions that would otherwise be lost due to sparse coverage or uneven CpG distribution.

## Intra-region variability

Optiles evaluates intra-region methylation variability by calculating the standard deviation of methylation values within each region across individual samples or biological replicates, using the *compute_beta_sd_regions* function. The input data consists in CpG-level methylation values together with their corresponding mapped genomic regions and specify the sample group across which the variability should be computed. In general, regional methylation summaries are typically computed as averages of CpG methylation levels within a region (Akalin et al. 2012), some methods weight this average by read coverage to account for uneven sequencing depth (Hansen, Langmead and Irizarry 2012, Feng, Conneely and Wu 2014). We believe that, beyond calculating a region average methylation value, examining the variability of individual CpG sites can provide insight into both the consistency of the regional signal and biologically relevant heterogeneity.

Figure 1 (C-D-E) illustrates how flagging this variability can provide an additional layer of insight. In Figure 1C, a single fully non-methylated CpG site drives the high variability in this otherwise highly methylated region. This could result from genuine lack of methylation or technical factors. In this case, the site carries a known single-nucleotide polymorphism (SNP) in which the guanine following the cytosine is altered, thereby eliminating the CpG site and preventing methylation. If polymorphisms are not filtered at the outset, this approach allows case-by-case identification of variants that need to be removed. While removing all sites overlapping with common polymorphisms a common practice, it may not be ideal, especially when dealing with single cell types where SNP lists might be incomplete or not readily available (LaBarre et al. 2019). By flagging such regions with inconsistent methylation, the user remains informed about potential bias without the need of completely removing all species-level variation. Such regions can yield skewed mean methylation values, and the variability flag highlights them as candidates for closer examination or possible exclusion from downstream analyses.

In contrast, Figure 1D illustrates heterogeneous methylation across the CpGs within the region, indicating a pattern that might reflect regulatory complexity. Notably, this pattern maps to a proximal enhancer-like signature, suggesting that such heterogeneity may be associated with regulatory activity, transcription factor binding variability, or context-specific gene expression. Recent work highlights that methylation heterogeneity can arise from the dynamic interplay between DNA methyltransferases (DNMT) and ten-eleven translocation (TET) mediated modification cycles, with intermediate patterns reflecting either gradual transitions or regulatory states stabilized by transcription factor binding (Shi et al. 2021). Such heterogeneity has been linked to context-specific gene expression and may provide more sensitive indicators of regulatory activity than average methylation levels alone (Lin et al. 2023). The flag highlights potentially meaningful heterogeneity that warrants further investigation; this variability cannot be generalized across the entire region and instead requires higher-resolution analysis to fully understand its functional implications.

In Figure 1E, the CpGs within the region display consistently similar methylation levels, indicating minimal intra-region variability. This region overlaps a CpG island (CpG43) located near the transcription start site (TSS) of the *ERLIN1* gene. The uniform demethylation pattern is consistent with the well-established tendency of CpG islands in promoter regions to remain unmethylated (Bird 2002, Deaton and Bird 2011). In such cases, summarizing the region with a single average methylation value is both appropriate and representative of its biological state.

This additional layer of information improves the interpretability of regional methylation patterns, especially in complex or heterogeneous datasets. By integrating intra-region variability with CpG density into a single scoring framework we define the InfoScore, computed calculated as the ratio between CpG density (number of CpGs divided by region length) and the standard deviation of methylation values within the region. With this score, OpTiles helps prioritizing regions that are both well-supported by data and internally consistent and additionally by flagging those with variability that could indicate underlying biological or technical factors. This enables more informed prioritization without prematurely excluding regions that may hold biological relevance.

## Additional functionalities

Beyond tile optimization and intra-region variability assessment, OpTiles includes a suite of functions to map user-defined regions to genomic annotations. This is achieved through integration with biomaRt database (Durinck et al. 2009), using the *biomart_annotation* function, which requires a specified database and dataset, and accepts optional filters such as chromosomes, attributes, attribute page, gene biotype(s), and promoter distance. The function enables annotation of methylation regions relative to known genomic features (e.g., promoters, genes, or CpG islands if annotation file is provided, as shown in the manual available on https://github.com/fhaive/OpTiles). A key aspect of this mapping functionality is the introduction of two overlap metrics that provide more nuanced control over region-feature associations. First, OpTiles computes the percentage of base-pair overlap between a given region and the target genomic feature. Second, it quantifies how many CpGs within the region fall inside the overlapping window. These metrics offer complementary views of region relevance and can be used as customizable filters to refine annotation outputs. This allows users to prioritize overlaps that are not only spatially extensive but also rich in informative CpGs, thereby supporting more tailored and interpretable downstream analyses.

## Conclusion

OpTiles extends the standard tiling-windows approach by refining region definitions, supporting data-driven prioritization, and enhancing interpretation of heterogeneous or complex datasets. To improve region definition in DNA methylation analysis, OpTiles implements a data-adaptive framework that considers the non-uniform nature of CpG distribution and sequencing coverage. By repositioning tiling windows to better reflect the actual distribution of sequenced CpGs, OpTiles preserves region size while increasing the retention of informative CpGs that might otherwise be excluded by genome fixed-window strategies. Beyond optimizing tile boundaries, OpTiles incorporates intra-region variability as an additional metric to assess the reliability and biological relevance of regional methylation summaries. Variability across CpGs within a region can arise from technical noise, cell-line specific SNPs, other sources of (epi)genetic variation, all of which are important to recognize. Rather than being treated as variation, OpTiles quantifies it alongside CpG density to offer a composite score, the InfoScore, designed to support informed region selection. This score helps to prioritize regions where methylation values are likely to be more stable and representative, and others that are instead more variable and inconsistent in methylation value that might need more careful evaluation. In parallel, the annotation and overlap-scoring functions provided by OpTiles offer researchers a finer-grained toolkit for contextualizing methylation patterns. This is especially valuable in studies aiming to link methylation changes to regulatory regions, where the extent and density of overlap carry different but complementary implications. Rather than replacing existing workflows, OpTiles is designed to complement and enhance them by introducing a datadriven, flexible approach to region definition and interpretation. Overall, it complements existing pipelines by refining region definition, enhancing data quality, and improving the biological relevance of epigenomic analyses.

## Data availability

The package, the manual and the example script are available at https://github.com/fhaive/OpTiles. The repository also contains links to documentation and installation instructions The data used for the analysis are available on Data are available on Zenodo at https://doi.org/10.5281/zenodo.16961293.

## Author contribution

**GM**: conceptualization, data curation, former analysis, investigation, methodology, software, validation, visualization, writing original draft, writing review & editing. **LM**: conceptualization, former analysis, methodology, supervision, writing original draft, writing review & editing. **GdG**: conceptualization, supervision, writing review & editing. **JM**: conceptualization, supervision, writing review & editing. **AF**: supervision, writing review & editing. **AS**: supervision, writing review & editing. **DG**: conceptualization, funding acquisition, project administration, resources, supervision, writing review & editing.

